# Mass spectrometric analysis and biostimulatory effects of boar seminal gel, saliva and semen in pigs

**DOI:** 10.1101/2023.02.08.527243

**Authors:** Sunil Kumar, Rafiqul Islam, Anesha Chanda, Pranab Jyoti Das, Santanu Banik, Keshab Barman, Seema Rani Pegu, Swaraj Rajkhowa, Vivek Kumar Gupta

## Abstract

The present study aimed to identify novel biostimulatory compounds in boar seminal gel (SG), saliva and semen using Gas chromatography-mass spectrometry (GC-MS). SG alone and its combined application with saliva (SG+saliva) and semen (SG+semen) also studied to train young boars. SG alone and SG+Saliva investigated for estrus induction in gilts and sows. Distilled water (DW) exposure was kept as control. SG, saliva and semen screened for total 105, 96 and 89 compounds. The highest concentration was of alkanes followed by sugar alcohols, then hydrocarbons, amino acids and fatty acids. Elaidic acid is the novel compound identified in pigs. Other compounds were tridecenol, undecane, hexadecone, eicosane, tetracosane etc. Further, young males (64.86%) were able to get trained. Significant higher (p<0.05) number of males got trained in exposure to SG (80%), SG+saliva (75%) and SG+semen (75%) than control. The time (hrs) taken by young boars to get trained on exposure to combination of SG+saliva (244±22.19) and SG+semen (216±13.14) was lesser (p<0.05) than SG (356±61.85) alone. Interval (hrs) for exhibition of different sexual behaviour by males on exposure to SG, saliva and semen was lesser (p<0.05) than control. Estrus was induced in 61.25% of females. Significant (p<0.05) higher number of females showed estrus response to exposure of SG (72.72%) and SG+saliva (69.23%) than control. Interval taken to exhibit estrus was lesser (p<0.05) in females exposed to SG+saliva (201.88±12.66) than SG (262.14±20.06) alone. Interval (hrs) for exhibition of different sexual behaviour by females on exposure to SG+saliva was lesser (p<0.05) than control. In conclusion, novel compounds with biostimulatory properties have been identified in boar SG, saliva and semen. The combined exposure of SG with saliva and semen have more intense biostimulation effect than SG alone. Such compounds and biostimulatory effects can be exploited for augmenting reproductive efficiency in pigs.

## INTRODUCTION

In the Livestock concern, biostimulation is a term coined to describe the stimulatory (positive) effects of a male on estrus, ovulation, or pregnancy (Chenoweth, 1983). Identification of pheromones, chemical signaling and their effects in insects are widely studied (Meinwald et al. 1974, Renou et al. 1981, Grant et al. 2003). However, only few reports are available in mammals particularly livestock. The 5α-androst-16-ene-3-one is the only identified compound used commercially in mammals. Among livestock, sheep, goat, and swine producers routinely utilize the effect of males (Thomson and Savege, 1978; Dyck, 1988) in management procedures to enhance reproductive performance. For example, the presence of a male hastens the onset of puberty in ewe lambs (Dyrmundsson and Lees, 1972) and gilts (Brooks and Cole, 1970; Kirkwood et al., 1981), and certainly advances the onset of estrus in mature ewes (Oldham et al., 1978; Pearce and Oldham, 1988), goats (Shelton, 1960) and lactating sows (Rowlinson and Bryant, 1974). However, boar parading for estrus detection or transferring females in mature boar shed for estrus induction or minimizing age at puberty is a difficult task due to problem in animal control. Moreover, it consumes a lot of time in shifting and transportation of animals and subsequent possibilities of injury of animals. All these facts make the conventional protocols too cumbersome for modern intensive farming. Further, the use of injectable hormones in females for estrus induction and synchronization has its limitations such as ease of availability, cost, skilled manpower and time. Such problem can be overcome by developing commercial applications like nasal spray, gel or ointment so that animal handling can be minimized and some non-invasive procedures may be employed instead of injectable ones. One such example in the pig is boar saliva which has pheromones namely 5α-androst-16-en-3-ol, 5α-androst-16-ene-3one and quinoline. The compounds 5α-androstenone or 3α-androstenol were earlier identified (Melrose *et al*. 1971) and is used commercially (Boar Mate; Antec. A. H. International, Ltd.) in limited countries and still has not got licensed to use across the globe. This provides an opportunity to identify new compounds in saliva, semen and seminal gel so that newer preparation can be prepared for enhancing reproductive efficiency in female pigs. Further, In the Artificial Insemination programme in pigs, training of males for semen collection is one of the toughest tasks as the dummy used is non-living, unlike cow and buffalo bulls where a live dummy is used to train the male for semen collection. Training the young boars requires patience, a good understanding of the psychological behaviour of the male, handler’s experience and comfort environment. However, many young males fail to get trained possibly because of a lack of natural biostimulation which usually they get in presence of their herd mates which may be either male or female. Most of the young males are culled or rejected because they are difficult to train for semen collection. Then such males are either used in natural service or culled considering probable reason of low or lack of libido. To mask natural biostimulation, seminal gel (SG) and saliva of already trained boars offers a good opportunity to train other young males. The hypothesis is supported by the evidence that an incoming second boar mounts with more interest and aggressively on the dummy on which a recent prior collection has already been made. This might be due to leftover salivary secretions, body smell and initial drops of semen from the previous boar on the dummy. Therefore, identification of novel pheromones would pave the way to develop preparation like nasal spray, ointment or gel for inducing or synchronizing estrus in sows and reducing the age at puberty in gilt effectively. Further, it can be used to enhance libido or improve semen quality in young boars. The best samples for such identification will be seminal gel, saliva and semen. Biochemical analysis of components in seminal gel secreted from boar semen has been reported with limited composition only. However, it is not characterized for active components having a biostimulatory effect that influences sexual responses. To the best of our knowledge, it is not known to date that boar seminal gel, semen and saliva (except 5α-androst-16-en-3-ol, 5α-androst-16-ene-3one and quinoline) contains compounds having pheromonal properties. Further, no reports are available on the use of semen, saliva and seminal gel for biostimulatory effects in young male and female (gilt and sow) pigs for estimating different sexual behavioural parameters. Therefore, the present investigation was planned for the identification of pheromonal compounds in boar’s seminal gel, semen and saliva along with the comparative estimation of sexual responses in response to biostimulatory effects in young boars, gilts and sows.

## MATERIALS AND METHODS

### Experimental animals and sample collection

The present study was conducted under intensive farm management conditions at Indian Council of Agricultural Research-National Research Centre on Pig, Rani (Assam), India. Semen was collected from adult healthy trained boars by double gloved hand method per standard protocol. The SG mass was separated over the collection flask topped with a filter during semen collection. The drooling saliva of boars was collected in sterile vials during semen collection. Initially, for GC-MS analysis, the gel mass, saliva and semen were kept immediately at −20°C for 24 hours in triplicate. For biostimulation applications, freshly collected gel mass (~15ml) and saliva (~10ml) were used for training of young males and estrus induction in females. Subsequently, for the training of males, young boars (n=37; age 8.67±0.10 months) were divided into four groups (Gp; three treatments and one control). In the treatment groups, a total of 31 males were exposed twice daily for 30 minutes for 15 days, to SG alone (Gp-I; n=15), and its combinations SG+saliva (Gp-II;n=8), SG+semen (Gp-III; n=8) by rubbing on the dummy. The interval (hrs) from the time of first exposure to exhibition of sexual behavioural parameters such as interest in the smell of treatments applied (SG, SG+saliva, SG+semen), interest in dummy, biting to dummy, salivation, licking to dummy, erection of the penis, mounting on dummy and first semen collection was estimated. Six young males (Gp-IV; n=6) were exposed to distilled water on a dummy as a control.

Finally, a total of 80 females (gilts-50; sows-30) were divided into three groups. Sows (2^nd^ - 3^rd^ Parity) with more than 10 days (26.93±0.14) of weaning and gilts more than 6 months (6.57±0.05) of age and 70.1±1.41 kg body weight along with history of absence of estrus expression were used as experimental animals. In group-A, females (Gp-A; n=55) were exposed to gel mass alone and in group- B, females (Gp-B; n=13) were exposed to combination gel mass and saliva. In group- C, females (Gp-C; n=12) were exposed to distilled water (DW) as control animals. The exposure was given twice daily 30 minutes for 15 days by keeping the exposure material in the pen of female near to snout as much as possible.After exposure, interval (hrs) from first exposure to exhibition of interest in smell, restlessness, urination, homosexual mounting, vulvar swelling, redness of vulva and positive back pressure were noted. The estrus was confirmed by back pressure test .

### Sample preparation for GC-MS

#### Gel

Frozen samples were brought to room temperature and 500 mg samples were added into 1mL of dichloromethane and kept aside for 24 hrs. After centrifuging the sample at 1200 rpm (4°C) for 10 min, supernatant was collected and injected (1μL) into GC-MS for analysis.

#### Saliva

samples were thawed and added 100μL of each sample into the eppendorf tube and then added 300μL of methanol. After that mixture was vortexed for 3 min and centrifuged at 12000 rpm (4°C) for 10 min. A volume of 200μL of supernatant was collected and evaporated under a vacuum concentrator. Then samples were derivatized by methoxylation followed by the silylation process. For this, dried extract was suspended in 90μL of methoxyamine hydrochloride in pyridine (20mg/mL), vortexed for 2 min and then heated in a dry bath at 60°C for 1.3hr. After that added 200μL of silylation reagent (N-Methyl-N-(trimethylsilyl) trifluoroacetamide +0.1% Trimethylchlorosilane) and heated in a dry bath at 60°C for 2hr. Then samples were evaporated under a vacuum concentrator and reconstituted with 100μL of n-hexane and injected (1μL) into GC-MS for sample analysis.

### Instrumentation

The dried sample after derivatization was mixed with n-hexane (300μL) and vortexed. An aliquot of 1 μL was injected into GC-MS (GC 9000 and MS of G7077B, Agilent Technologies, Palo Alto, CA, USA) for further analysis. The GC-MS analysis was carried out with the following conditions. The injector temperature was maintained at 280°C and the oven temperature program with initial oven temperature was held at 70°C for 4 min, then increased to 300 °C at a rate of 10°C/min and finally held for 5 min. Helium was a carrier gas with a flow rate of 1 ml/min. The DB - 5MS capillary column (30 m X 250 μm i.d. X 0.25 μm film thickness in splitless mode was used. The data was acquired with electron ionization mode at 70 eV. The MS source and transfer line temperature were set at 230°C and 290°C respectively. Full scan mass spectra were acquired in the mass range of m/z 29-600 with an initial solvent delay of 6 min.

### Data analysis

The peaks acquired by GC-MS were identified by comparing their mass spectra with NIST library mass spectra’s were selected for the compounds having more than 70% of matching score in NIST library. In metabolite analysis, the enrichment ratio for a particular pathway was defined as the ratio of measured masses that map to metabolites within the pathway to its size. For biostimulation studies, the group difference for sexual behavioural parameters was estimated by using student’s t test. Data was analyzed using statistical package version 16.0 (SPSS Inc., Chicago, IL, USA).

## RESULTS

### Mass spectrometric analysis of boar seminal gel, saliva and semen

The boar seminal gel, saliva, and semen were subjected to chemical analysis by using optimized GC-MS parameters and the resultant gas chromatograms for seminal gel (a), saliva (b) and semen (c) are presented in Fig 1. The compositions of the chemical compounds present in boar seminal gel, saliva, and semen identified by GC-MS analysis with their retention time (RT), molecular formula, molecular weight and area (%) are presented in Table 1. The identified compounds were subjected to metabolite analysis using MetaboAnalyst 5.0 and it was found that the identified compound belongs to categories of alkanes, hydrocarbons, amino acids, sugar alcohols, unsaturated fatty acids, branched fatty acids and fatty alcohols. The enrichment ratio (Fig 2.) indicated that the highest concentration was of alkanes followed by sugar alcohols, then hydrocarbons, amino acids and fatty acids.

**FIGURE 1.**
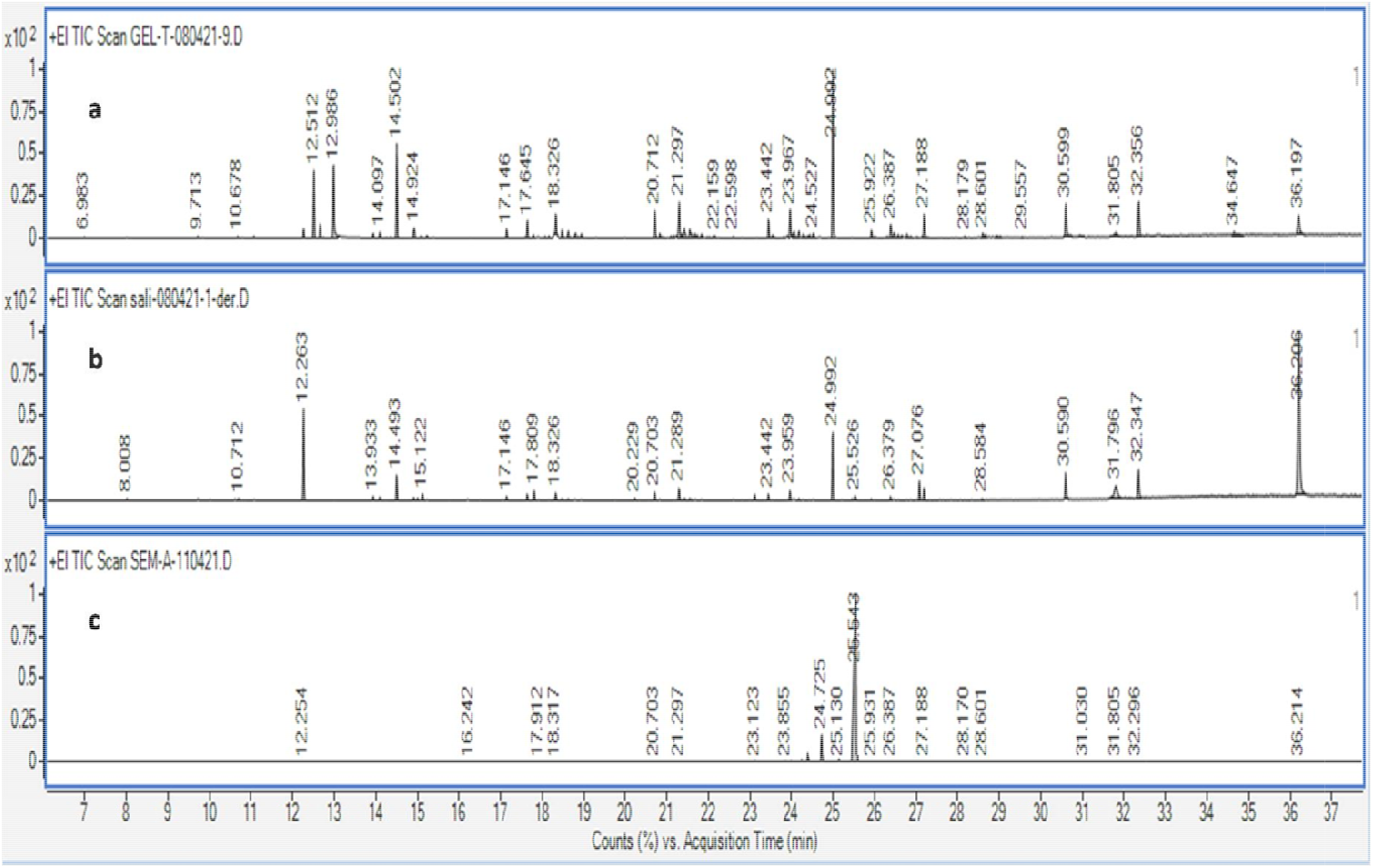
GC-MS chromatograms (count vs acquisition time) of boar seminal gel (a), saliva (b) and semen (c). Acquisition time (minutes) on X-axis and percentage count or intensity on Y-axis. The x-axis of the gas chromatogram shows the amount of time taken for the analytes to pass through the column and reach the mass spectrometer detector. The peaks that are shown correspond to the time at which each of the components reached the detector. In the chromatogram, the area is based on the number of counts taken by the mass spectrometer detector at the point of retention. The figure depicted is extracted ion chromatogram (s) for SG (a), saliva (b) and semen (c) for a particular m/z value where m stands for mass and z stands for charge number of ions for detected compounds.

**Table 1.** Compounds identified in boar seminal gel, saliva and semen using GC-MS analysis^#^

| SN | Compound | Mwt | M.F | RT | Area(%) | Score |
| --- | --- | --- | --- | --- | --- | --- |
| **a** | **Seminal Gel** |  |  |  |  |  |
| 1 | Nonadecane | 268 | C<sub>19</sub>H<sub>40</sub> | 6.933 | 0.46 | 96.49 |
| 2 | Tridecenol | 198 | C<sub>13</sub>H<sub>26</sub>O | 10.678 | 0.09 | 96.81 |
| 3 | 9-(E)-octadecenoic acid | 282 | C<sub>18</sub>H<sub>34</sub>O<sub>2</sub> | 12.263 | 1.52 | 95.20 |
| 4 | Undecane | 156 | C<sub>11</sub>H<sub>24</sub> | 12.996 | 56.23 | 95.18 |
| 5 | 4-heptanol-2-methyl | 130 | C<sub>8</sub>H<sub>18</sub>O | 14.097 | 0.89 | 80.04 |
| 6 | 2-isopropyl-5-methyl-1-heptanol | 172 | C<sub>11</sub>H<sub>24</sub>O | 17.146 | 1.98 | 94.90 |
| 7 | Hexadecane | 226 | C<sub>16</sub>H<sub>34</sub> | 17.645 | 2.01 | 71.24 |
| 8 | 3-ethyl-3-methyl heptane | 142 | C<sub>10</sub>H<sub>22</sub> | 20.712 | 5.23 | 75.68 |
| 9 | Tetracosane | 338 | C<sub>24</sub>H<sub>50</sub> | 22.199 | 0.24 | 86.65 |
| 10 | Henicosane | 296 | C<sub>21</sub>H<sub>44</sub> | 23.442 | 2.98 | 85.08 |
| 11 | Dodecane | 170 | C<sub>12</sub>H<sub>26</sub> | 24.557 | 0.89 | 58.95 |
| 12 | Nonane | 128 | C<sub>9</sub>H<sub>20</sub> | 24.992 | 100 | 71.35 |
| 13 | 2,3-butanediol | 90 | C<sub>4</sub>H<sub>10</sub>O<sub>2</sub> | 27.188 | 3.25 | 95.20 |
| 14 | 2-methyl butyric acid | 102 | C<sub>5</sub>H<sub>10</sub>O<sub>2</sub> | 28.601 | 1.23 | 95.18 |
| 15 | 3-isopropyl-5-methyl-2-hexanone | 156 | C<sub>10</sub>H<sub>20</sub>O | 29.557 | 0.09 | 91.40 |
| 16 | 2-furanol | 84 | C<sub>4</sub>H<sub>4</sub>O<sub>2</sub> | 30.995 | 6.23 | 51.92 |
| 17 | 2,6 dimethyl decane | 170 | C<sub>12</sub>H<sub>26</sub> | 31.805 | 0.51 | 89.19 |
| 18 | 3,4-hexanedione | 114 | C<sub>6</sub>H<sub>10</sub>O<sub>2</sub> | 32.356 | 6.89 | 88.00 |
| 19 | 3-isopropyl-6,10-dimethyl undecane-2-ol | 242 | C<sub>16</sub>H<sub>34</sub>O | 34.647 | 0.08 | 82.94 |
| 20 | 2-bromotetradecane | 277 | C<sub>14</sub>H<sub>29</sub>Br | 36.197 | 4.56 | 95.14 |
| **b** | **Saliva** |  |  |  |  |  |
| 1 | Glycerol | 92 | C<sub>3</sub>H<sub>8</sub>O<sub>3</sub> | 8.008 | 0.25 | 90.38 |
| 2 | DL-alanine | 89 | C<sub>3</sub>H<sub>7</sub>NO<sub>2</sub> | 10.712 | 0.51 | 95.08 |
| 3 | 4-bromobutyric acid | 167 | C<sub>4</sub>H<sub>7</sub>BrO<sub>2</sub> | 12.263 | 45.26 | 96.60 |
| 4 | Undecane | 156 | C<sub>11</sub>H<sub>24</sub> | 15.122 | 1.89 | 95.08 |
| 5 | 5-chloro-thiopene-2-carbonyl chloride | 181 | C<sub>5</sub>H<sub>2</sub>Cl<sub>2</sub>OS | 17.146 | 1.34 | 96.49 |
| 6 | 6-fluoro-2-trifluoromethyl benzoic acid | 360 | C<sub>20</sub>H<sub>12</sub>F<sub>4</sub>O<sub>2</sub> | 17.809 | 1.95 | 80.04 |
| 7 | Dodecane | 170 | C<sub>12</sub>H<sub>26</sub> | 18.326 | 1.38 | 94.90 |
| 8 | D-fructose | 180 | C<sub>6</sub>H<sub>12</sub>O<sub>6</sub> | 20.229 | 0.76 | 71.24 |
| 9 | Hexadecane | 226 | C<sub>16</sub>H<sub>34</sub> | 20.703 | 1.23 | 95.48 |
| 10 | Chiro-inositol | 180 | C<sub>6</sub>H<sub>12</sub>O<sub>6</sub> | 21.289 | 2.12 | 88.13 |
| 11 | 3-ethyl-3-methyl heptane | 142 | C<sub>10</sub>H<sub>22</sub> | 23.959 | 1.82 | 70.43 |
| 12 | Cyclobutylamine | 71 | C<sub>4</sub>H<sub>9</sub>N | 24.992 | 18.96 | 83.15 |
| 13 | Androst-16 en-3-ol | 274 | C<sub>19</sub>H<sub>30</sub>O | 25.526 | 1.02 | 80.11 |
| 14 | 1-octanol | 130 | C<sub>8</sub>H<sub>18</sub>O | 27.076 | 3.26 | 73.63 |
| 15 | 3-hexanone | 100 | C<sub>6</sub>H<sub>12</sub>O | 28.584 | 0.09 | 83.23 |
| **c** | **Semen** |  |  |  |  |  |
| 1 | D-penitol | 194 | C<sub>7</sub>H<sub>14</sub>O<sub>6</sub> | 12.254 | 0.09 | 77.99 |
| 2 | Methyl-alpha-glucofuranoside | 194 | C<sub>7</sub>H<sub>14</sub>O<sub>6</sub> | 16.242 | 0.11 | 84.70 |
| 3 | beta-D-galactofuranose | 180 | C<sub>6</sub>H<sub>12</sub>O<sub>6</sub> | 17.912 | 0.06 | 92.94 |
| 4 | Tetradecane | 198 | C<sub>14</sub>H<sub>30</sub> | 20.703 | 0.04 | 82.69 |
| 5 | Ethylenimine | 43 | C<sub>2</sub>H<sub>5</sub>N | 23.123 | 0.09 | 63.99 |
| 6 | L-proline | 115 | C<sub>5</sub>H<sub>9</sub>NO<sub>2</sub> | 23.855 | 0.02 | 84.27 |
| 7 | Alanine | 89 | C<sub>3</sub>H<sub>7</sub>NO<sub>2</sub> | 24.268 | 2.56 | 96.14 |
| 8 | L-glutamic acid | 147 | C<sub>5</sub>H<sub>9</sub>NO<sub>4</sub> | 24.725 | 19.26 | 95.08 |
| 9 | 3-hexanone | 100 | C<sub>6</sub>H<sub>12</sub>O | 25.130 | 0.13 | 96.49 |
| 10 | Undecane | 156 | C<sub>11</sub>H<sub>24</sub> | 25.543 | 100 | 90.42 |
| 11 | 3-heptanone | 114 | C<sub>7</sub>H<sub>14</sub>O | 26.387 | 0.08 | 96.81 |
| 12 | Octane | 114 | C<sub>8</sub>H<sub>18</sub> | 27.188 | 0.03 | 70.76 |
| 13 | Dodecane | 170 | C<sub>12</sub>H<sub>26</sub> | 27.559 | 0.12 | 88.95 |
| 14 | Hexadecane | 226 | C<sub>16</sub>H<sub>34</sub> | 28.170 | 0.05 | 83.23 |
| 15 | 2-bromo dodecane | 249 | C<sub>12</sub>H<sub>25</sub>Br | 28.601 | 0.07 | 71.30 |
| 16 | 1-butene | 56 | C<sub>4</sub>H<sub>8</sub> | 31.030 | 0.04 | 77.99 |
| 17 | Eicosane | 282 | C<sub>20</sub>H<sub>42</sub> | 31.805 | 0.09 | 85.68 |
| 18 | Tetracosane | 338 | C<sub>24</sub>H<sub>50</sub> | 32.296 | 0.8 | 86.65 |
| 19 | 2-propanol | 60 | C<sub>3</sub>H<sub>8</sub>O | 36.241 | 0.6 | 91.25 |

**FIGURE 2.**
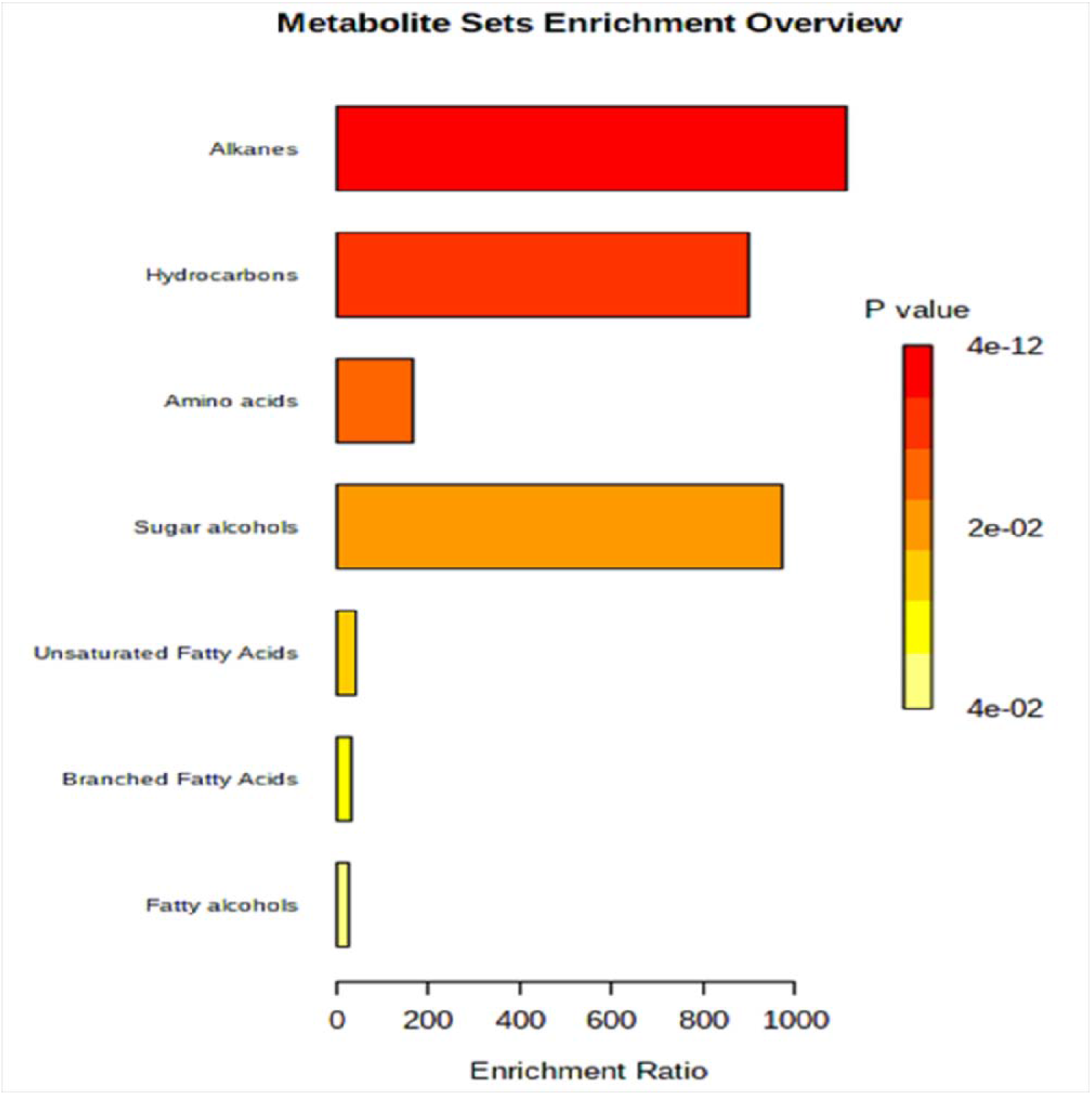
The enrichment ratio of identified compounds in boar seminal, gel and saliva. The enrichment ratio for a particular pathway was defined as the ratio of measured masses that map to metabolites within the pathway to its size.

**FIGURE 3.**
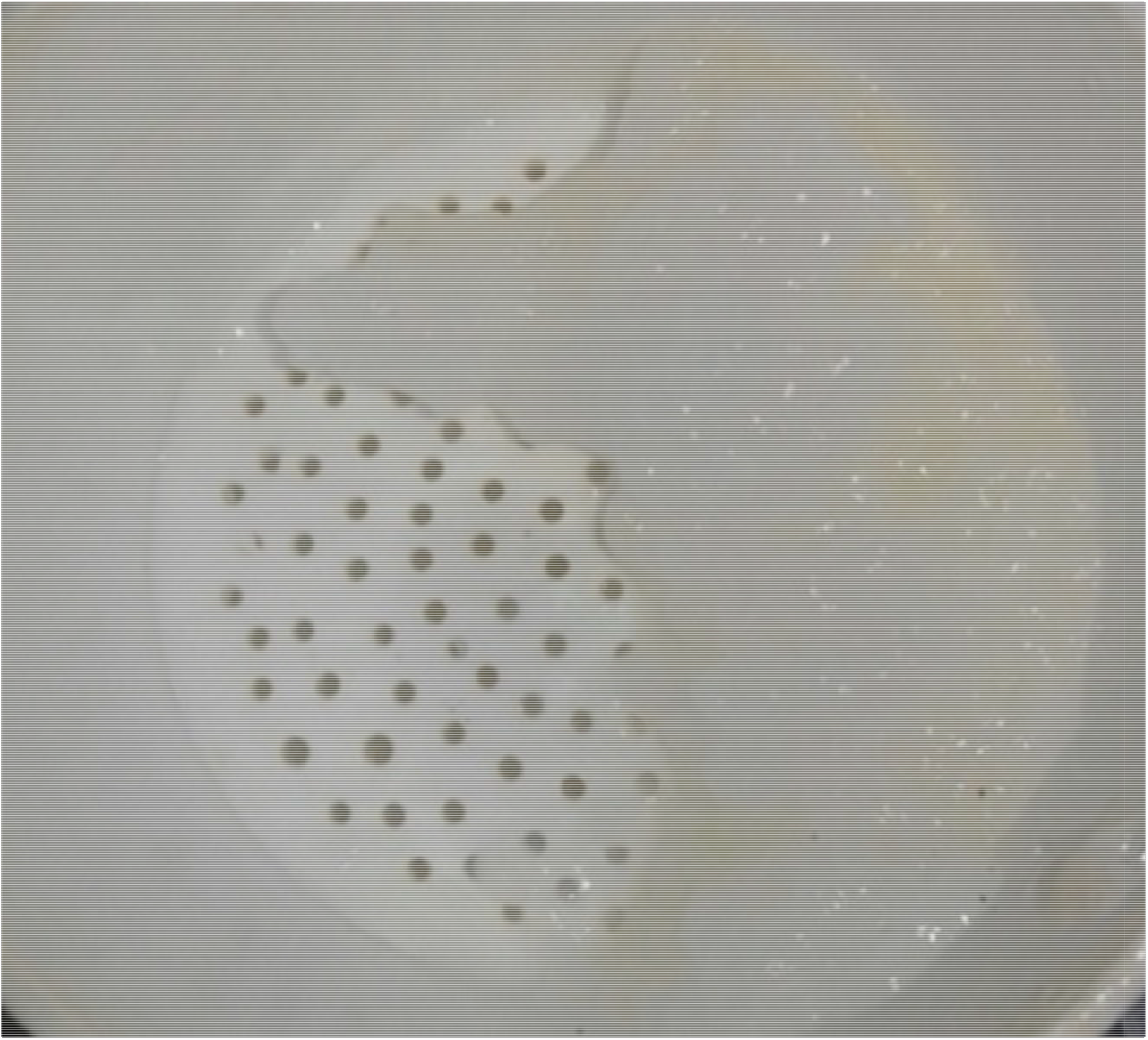
The physical form of boar seminal gel may be due to the presence of heneicosane. The gel like consistency of boar seminal gel may be due to heneicosane as observed in freshly collected boar seminal gel which is quite similar to physical form.

#### Seminal Gel

On GC-MS Analysis, a total of 105 compounds were identified in boar seminal gel. Out of which, 30 compounds were found to have pheromonal properties. Among 30 compounds, 12 were found to be potential pheromones as per literature. The compounds identified are namely as Non-decane, Tridecenol, 9-octadecenoic acid, Undecane, 4-heptane-2-methyl, 2-isopropyl 5-methyl-1-heptanol, Hexadecone, 3-ethyl-3 methyl heptone, Tetracosane, Henicosane, Dodecane, Nonane, 2,3-butaanediol, 2-methyl butyric acid, 3-isopropyl 5-methyl-1-heptol, 2-furanol, Hexadecone, 2,6 dimethyl decone, 3,4 hexanedione,3-isopropyl-6,10 dimethyl undecane-2-ol, 2-bromotetradecane, Pyruvic acid, Alanine, L-norvaline, Alanylalanine, L-glutamic acid, DL_alanine, Decane, D-mannose, α,D-glucopyranoside.

#### Saliva

A total of 96 compounds were screened from boar saliva. Compounds identified with probable pheromonal properties were 21, where six are expected to be potential pheromones including Androst-en-ol. The identified compounds included Glycerol, DL-alanine, 4-bromobutyric acid, Undecane, 5-chloro-thiopene-2 carbonyl chloride, 6-flouro-2trifluro methyl benzoic acid, Dodecane, D-fructose, Hexadecane, Chizo-inositol, 1,2,6-isopylidene-α-D-glucofuranose, 3-ethyl-3 methyl heptanes, Cyclobutylaine, Androst-16 en-3-ol, 1-octanol, 3-hexanone, Decane-3 ethyl-3methyl, Eicosane, 2-pentadecanyl, 3ethyl-3methyl heptone and Henicosane.

#### Semen

Total 89 compounds were screened from boar semen out of which 27 compounds identified as pheromonal nature and seven identified as potential pheromones. The identified pheromonal compounds include D-pemitol, 3-hexane, Methyl α-lynofuranoside, β-D-galactofuranose, Tetradecane, Ethylenimine, L-proline, Alanine, L-glutamic acid, 3-hexanone, Undecane, 3-heptanone, Octane, Dodecane, Hexadecane, 2-bromo dodecane, 1-butene, Eicosane, Tetracosane, 2-propanaol, 3-hexanone, and 2,2,5,5,-tetramethyl-3-hexanone.

### Estimation of biostimulatory effect of seminal gel, saliva and semen for the training of young boars

The SG and its combination of saliva and semen were used in control with DW for the training of young males. The results for training of boar using SG alone and combined application with saliva and semen are represented using Table 2 and Fig 4. Out of total of 37 males, 24 (64.86%) males were able to get trained. However, among the treatment groups I, II and III, 77.41 % (24/31) was the success rate while the rejection rate (7/31) was 22.58 %. In Gp-I, it was observed that boars get trained (12/15; 80%) after 356±61.85 hrs while the corresponding interval (hrs) in Gp-II (6/8;75%) and Gp-III (6/8;75%) was 244±22.19 and 216±13.14 hrs respectively. Boars (n=6) exposed to DW (group-IV) were not able to get trained.

**Table 2.** Comparison of interval (hrs) from the time of first exposure to the observation of sexual behaviour parameters in boar training using seminal gel alone (I) and combination of with saliva (II) and semen (III) in comparison to distilled water as the control group (IV). The values with different superscripts (a, b and c) in different groups differ significantly (p<0.05)

| Group | Hours (hrs) from treatment/exposure of gel and its combination to the observation of sexual behaviour |  |  |  |  |  |  |  |
| --- | --- | --- | --- | --- | --- | --- | --- | --- |
|  | Smell to gel | Interest in Dummy | Biting to Dummy | Salivation | Licking to Dummy | Erection of Penis | Mounting on Dummy | Exposure to collection (Hrs) |
| I | 44±9.03<sup>a</sup> | 124±24.49<sup>a</sup> | 116±15.94<sup>a</sup> | 60±6.57<sup>a</sup> | 92±9.03<sup>a</sup> | 276±79.14<sup>a</sup> | 308±74.90<sup>a</sup> | 356±61.85<sup>a</sup> |
| II | 24±00 | 88±24.26<sup>b</sup> | 80±22.62<sup>b</sup> | 56±07.15 | 64±07.15<sup>b</sup> | 168±17.52<sup>b</sup> | 184±20.86<sup>b</sup> | 244±22.19<sup>b</sup> |
| III | 24±00 | 56±9.79<sup>c</sup> | 80±6.19<sup>b</sup> | 40±6.19<sup>b</sup> | 60±6.57 | 76±9.03<sup>c</sup> | 172±11.79<sup>b</sup> | 216±13.14<sup>bc</sup> |
| IV | 80±5.05<sup>b</sup> | 189.33±14.11<sup>d</sup> | 132±6.92<sup>c</sup> | 124.80±13.99<sup>c</sup> | 112.31±5.65<sup>c</sup> | 240±00<sup>d</sup> | Not mounted | Not trained |

**FIGURE 4.**
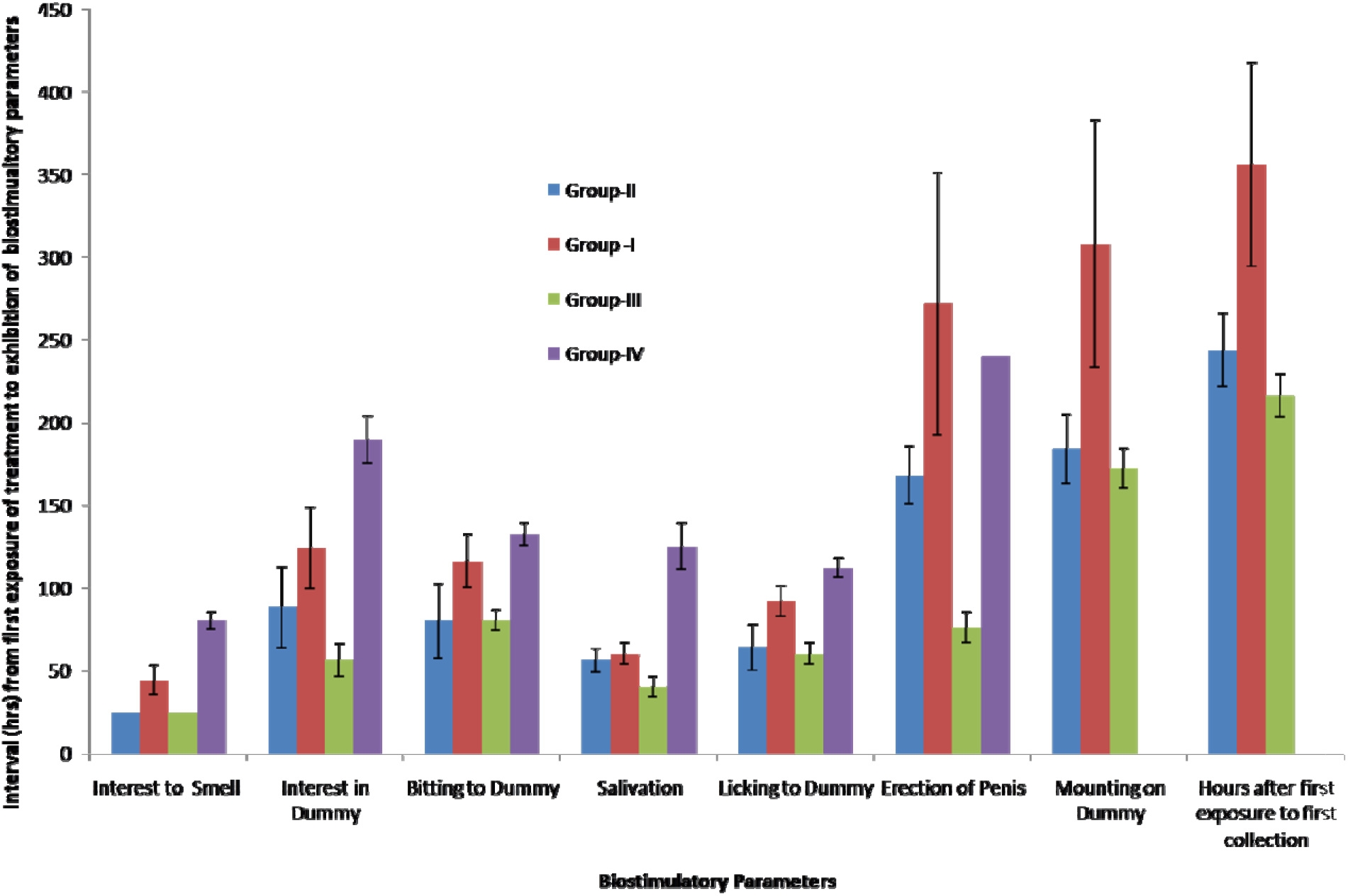
Comparison of effects on sexual behaviour for boar training using seminal gel alone and its combined application with saliva and semen. X-axis indicates the different sexual responses observed due to biostimulatory effects of different treatments such as seminal gel alone (Gp-1), SG+Saliva (Gp-II), SG+Semen (Gp-III) and distilled water (Gp-IV). Y-axis indicates the interval (hrs) for biostimualtory parameters exhibition (hrs) from the time of first exposure of different treatments.

### Estimation of biostimulatory effect of seminal gel and its combination with saliva for estrus induction in gilts and sows

The SG mass and its combination with saliva were used for oestrus induction in gilts and sows. Hormonal therapy and male introduction to males in females have their limitations as described in the introduction of the present study. The results for estrus induction using SG alone and combined application with saliva are represented using Table 3 and Fig 5. A significant (p<0.05) difference observed that 61.25% (49/80) females in total showed estrus while in Gp-A 72.72 % (40/55), Gp-B 69.23% (9/13) and in control Gp-C none of the female (0/12) showed positive back pressure. It was found that oestrus was observed in total 66.00% of gilts (33/50) and 53.33% sows (16/30) in the present study. However, in comparison to gilts of control Gp-C (0;0/6), higher percentage of estrus response was observed in gilts of Gp-A (75.67%;28/37) and Gp-B (71.42%;5/7). The corresponding results in sows were 66.66% (12/18), 66.66% (4/6), 0(0/6) in Gp-A, Gp-B and Gp-C respectively. Hence it is indicated that a significant (p<0.05) estrus induction response was observed in gilts and sows in the treatment groups than control animals. The positive back pressure test in all estrus induced females (n=40) was observed after 262.14±20.06 hrs from the time of first exposure in Gp-A. In Gp-B, females showed estrus response and the positive back pressure was observed after 216±12.64 hrs from the time of first exposure to a combination of SG+ saliva. Hence, it is indicated that the time taken to get in the heat was lesser (p<0.05) in Gp-B than Gp-A. Further, sexual behavioural parameters such as interest in gel, movement of tail, restlessness, frequent urination, redness of vulva, swelling of the vulva and mounting on pen mates were also observed for estrus observation in response to application of SG and saliva.

**Table 3.** Comparison on the interval (hrs) from the time of first exposure to observation of different sexual behavioural parameters by females on exposure to boar seminal gel (A), SG+saliva (B) and distilled water (DW). The values with different superscripts (a,b and c) between different groups differs significantly (p<0.05)

| Sexual behavioural parameters | Hours from first exposure of Gel alone, Seminal Gel+saliva and DW |  |  |
| --- | --- | --- | --- |
|  | Group A | Group B | Group C |
| Interest in smell | 28.20±1.92<sup>a</sup> | 26.11±2.11 | 118±6.60<sup>b</sup> |
| Movement of tail | 105.23±09.74<sup>a</sup> | 122.82±13.82 | 134±5.25 |
| Restlessness | 74.40±7.52<sup>a</sup> | 66.35±10.78 | 182±16.84 |
| Frequent Urination | 96±9.68<sup>a</sup> | 120±7.58<sup>b</sup> | 204±10.39<sup>c</sup> |
| Redness of Vulva | 78.6±6.18 | 79.05±11.92 | Not observed |
| Swelling of Vulva | 106.66±8.32 | 101.64±14.29 | Not observed |
| Mounting on Mates | 226.88±29.45<sup>a</sup> | 166.28±13.11<sup>b</sup> | Not observed |
| Positive Back Pressure | 262.14±20.06<sup>a</sup> | 201.88±12.66<sup>b</sup> | Not observed |

**FIGURE 5.**
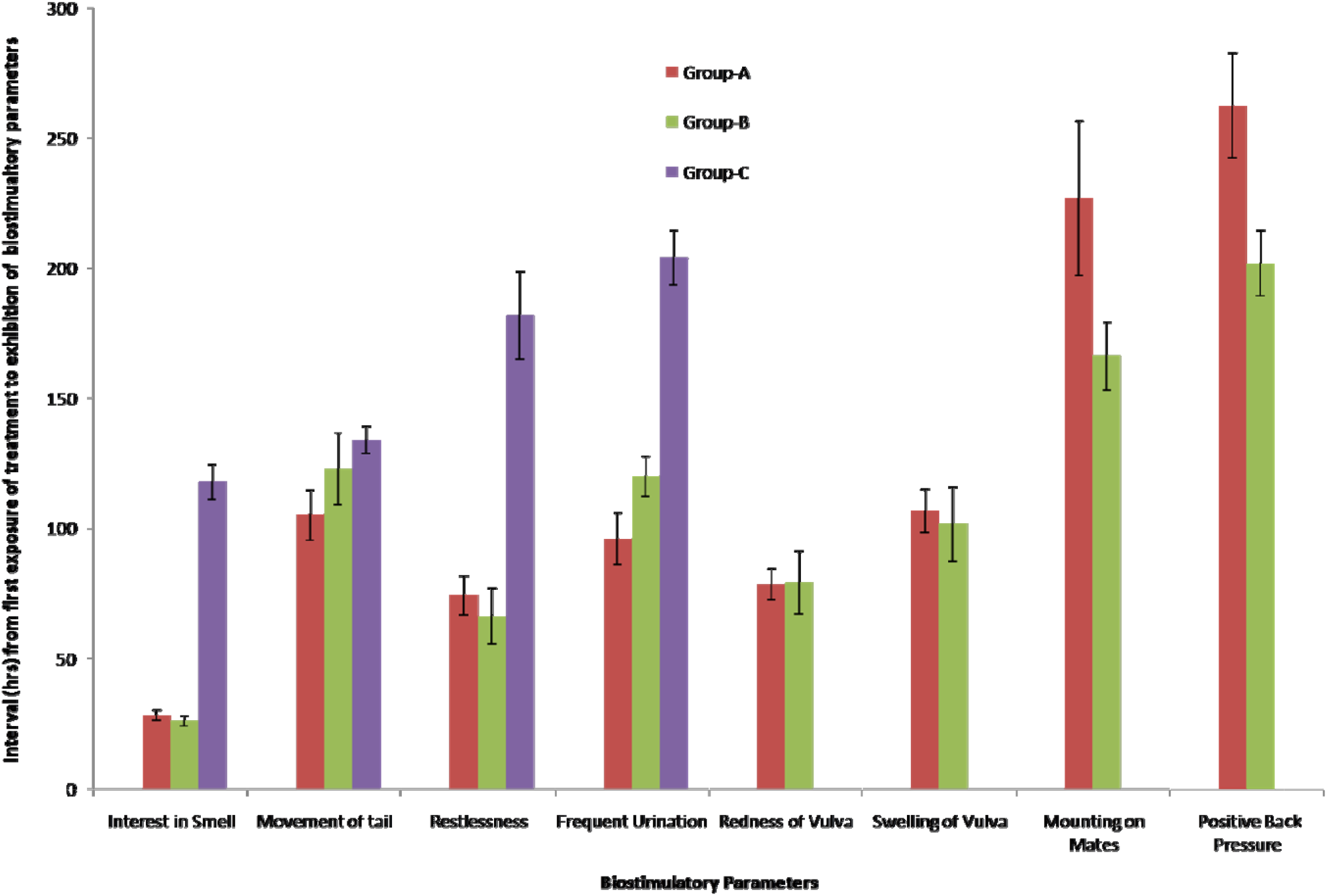
Comparison of effects on observations of different signs of estrus using seminal gel alone and combination of gel and saliva. X-axis indicates the different sexual responses observed du to biostimulatory effects by different treatments such as seminal gel alone (Gp-A), SG+Saliva (Gp-B) and distilled water (Gp-C). Y-axis indicates the interval (hrs) for biostimualtory parameters exhibition (hrs) from the time of first exposure of different treatments.

## DISCUSSION

Pheromones and other allelomimetic signals have proven effects on reproductive activity of domestic animals and can be used as control systems technologies to augment reproductive performance. The knowledge acquired on the effectiveness of biostimulation; the factor which conditions it and the biological mechanism which produces it in livestock species, allows its use as a reproductive management tool (Rekwot et al., 2001). The biostimulation technique offers a potentially useful and practical way to improve reproductive efficiency in livestock species in the tropics. Identification of pheromones or biostimulatory compounds required chemical analysis of source or site of production of compound. It is sometimes argued that mammals do not have many releaser pheromones that elicit a particular behaviour. However, some have been identified and more are likely to be discovered. One example is the ‘standing behaviour’ of an oestrous pig (*Sus scrofa*) female in response to the steroid sex pheromones, 3α-androstenol and 5α-androstenone, of the male pig in his saliva. The 5α-androstenone is the only mammalian pheromone to be synthesized and sold as a means of improving reproductive efficiency in livestock operations. However, it is the requirement to explore newer compounds or chemical signals which can provide further opportunities to explore the pheromonal and biostimulation biology in pigs. Therefore, seminal gel and semen were explored apart from the saliva, for identification of probable pheromones. Subsequently, these biological specimens were tested for estimating biostimulation for different sexual behaviours in male and female pigs.

GC-MS analysis of components in seminal gel secreted with boar semen has been done with respect to composition (Boursnell et al., 1970; Boursnell and Butler, 1973) only. It needs further exploration and validation for active ingredients having biostimulatory effect in pig. The only report (Takahashi et al., 2014) for the biochemical composition of boar seminal gel showed that SG contains O-glycans, galactosamine, O-glycosylated glycoprotein, tyrosine-protein kinase ZAP-70, mucin-like glycoproteins carrying heavily sialylated O-glycans. In the present investigation, most of the identified compounds have well-proven role as pheromone in insect or chemical ecology. These compounds are reported for the first time in swine. Such six compounds have been identified and isolated in the hair-pencil extracts of male *Acrolepiopsis assectella* (Thibout et al., 1994) by techniques of GC-MS. These compounds are n-alkanes: hexadecane (C16), heptadecane (C17), octadecane (C18), nonadecane (C19), eicosane (C20), and heneicosane (C21). Further, compound such as elaidic acid is patented in livestock and have a well-proven role in biostmulation in buffaloes. Further, 9(E)-Octadecenoic acid is a monounsaturated fatty acid and found to occur only during estrus in urine in cattle (Rajanarayanan and Archunan 2011), comes from vaginal fluid (Karthikeyan and Archunan 2013) and reported as biostimulant in bulls (Rajanarayanan and Archunan, 2011; Kekan et al., 2017). This compound enhances sperm production after exposure to the nasal region and this has been awarded a patent (Archunan and Rajanarayanan. 2010). This is the compound that can provide a breakthrough in the reproductive biology of swine. This compound is identified for the first time in swine species.

From the present study, heneicosane may be responsible for the gel like nature of the seminal gel. The freshly collected boar seminal gel (Fig.3) looks like to the organic form of heneicosane (Wikimedia, 2014). Further, it is a pheromone in queen or king *Reticulitermes flavipes* termites and encodes “royal status” (Funaro et al., 2017). It is an oviposition-attractant pheromone of larval origin in *Aedes aegypti* mosquito (Mendki et al., 2000). Undecane is an alarm pheromone of ant *C. obscuripes* (Fujiwara et al., 2006) and “Real” sex pheromone of *Formica lugubris*. Also, it is an oestrogen-dependent urinary sex pheromone of female mice (Achiraman et al., 2010). It is reported as an aggregation pheromone in *Blaberus craniifer* cockroaches and *Reticulitermes speratus* termites (Mitaka et al., 2020). It is an alarm pheromone component in *Camponotus obscuripes* ants (Mizunami et al., 2010). Tetracosane is a pheromone in Diaphanes (Wattanachaiyingcharoen et al., 2020), *Diuraphis noxia* (Bergman et al., 1990), *Lymantriidae* (Byers, 2006), Drosophila (Farine et al., 2012) and parasitoid wasps (Würf, et al., 2020). Dodecane has been reported as pheromone (Minaeimoghadam et al., 2017) and attracts gravid females to the larvae (Wittko, 1999). It is also a pheromone of rice stem borer (Liu et al., 2020), cerambycid beetle *Plagionotus detrit spp*. (Molander et al., 2018), male *Anelaphus inflaticollis* (Ray et al., 2009) and Tragosoma (Ray et al., 2012). Tridecenol is similar to Eicosen-1-ol (Pickett et al., 1982). In *Pityogenes chalcographus*, nonane is a component of the male aggregation pheromone. 2-methylbutyric acid is an insect pheromone precursor (Mittersteiner et al., 2017). It is a sex attractant pheromone blend of female house mice (Varner et al., 2018). Hexadecane is a component of pheromone of moth *Acrolepiopsis assectella* (Wittko, 1999) and clearwing moth, *Paranthrene diaphana* (Minaeimoghadam et al., 2017). The compound 2-bromotetradecan is reported as a sex attractant pheromone of house fly (Carlson et al., 1971; Yadav and Tissot, 1984). Eicosane has been reported as sex pheromones: *Acrolepiopsis assectella* (Renou et al. 1981), *Amauris niavius* (Meinwald et al., 1974) and *Orgyia leucostigma* (Grant et al., 2003).

In domestic animals, the chemical characteristics of these signaling pheromones and their metabolic pathways are still unclear. Little is known about how the production of signaling pheromones is influenced by hormonal status except for 5α-androstenone and 3α-androstenol. In some cases, the site of production of signaling pheromones is unknown. More information is needed in these areas to facilitate the practical use of signaling pheromones to improve the reproductive efficiency of domestic animals particularly pig.

For the training of young boars, a significant difference (p<0.05) for the training of males was observed between treatment groups than the control. Further, the interval (hrs) for different sexual behaviour parameters were also observed earlier in exposure to a combination of SG + semen followed SG + saliva and SG alone. It indicates that young boars in Gp-III, took a significant (p<0.05) lesser time to get trained followed by Gp-II and Gp-I. The earlier response in exposure to combinations than SG alone may be due to more quantity and potency of chemical signals produced by combination than in comparison to SG alone. The late or feeble response in the exposure to SG only may be due to fact the there might be positive interactions between the identified compounds between gel, saliva and semen which might be lacking in SG alone. Young boars exposed to DW showed sexual behavioural parameters such as smell to the water, interest in dummy, biting to dummy, salivation, licking to dummy and erection of penis with greater interval than as observed in exposure to SG and its combinations. These control grouped boars were unable to get trained; however, the erection of penis and interest in dummy was observed in the group. The inability to get trained in the control group may be due to a lack of biostimulation where only distilled water was used. From the present study, it is drawn that exposure to SG and its combination with saliva and semen provides biostimulation and is effective to train young boars. Thus, these biological materials can be used to train the males in artificial insemination programmes. However, it should be kept in mind that such biologicals should be from disease-free animals to prevent the disease transmission in the herd. For prevention of the diseases along with easy and wider use of such biological materials, it is advisable to use the identified compounds in purified form for the training of young males. Such practice will offer a breakthrough in the A.I. program in pigs. One such commercial preparation used in cow bulls in the name of Pherobull ™ (Patent INRA / UNCEIA; IMV Technologies, France) which is used as a nasal spray to stimulate sexual function and increase the reproductive capacity of the bull. This spray is made of synthetic molecules naturally occurring in the urine of cows in oestrus. Further, two compounds namely 4 methyl phenol and trans-verbenol were detected as reliable indicators of estrus in faeces of buffaloes (Karthikeyan et al., 2013). Hence, the present study, offers newer opportunities to prepare newer commercial preparations from SG, saliva and semen using identified probable pheromones.

For the purpose of estrus induction in gilts and sows, it was found that these sexual parameters were observed earlier on exposure in Gp-B than Gp-A. This might be due to intensive pheromonal effects of the identified compound in saliva and SG derived from the male counterpart. The introduction of males to reduce the age at puberty in gilts is well known. Similar findings reported that biostimulation of Sahiwal heifers with bull exposure resulted in greater expression of oestrous behaviour than in non-exposed heifers (Choudhary and Kamboj, 2019). It is reported that bull-cow biostimulation influences the resumption of ovarian activity of zebu (*Bos indicus*) cattle (Rekwot et al., 2000) following parturition possibly through olfactory cues (pheromones). The basis of all these reports relies on the chemical signaling of pheromonal compounds responsible for biostimulation in females. Hence, as per the present study, the use of such identified compounds from SG and saliva in chemical forms as a nasal spray, ointment or gel offers a new opportunity in pig estrus biology and it will help to make the estrus induction and synchronization easier, cheaper, economic and simple in pig farms. Such applications can be used by unskilled or semi-skilled persons in farms, in contrast, to the use of hormonal therapy where trained and skilled personnel is required. It will provide newer methods of non-invasive estrus induction and synchronization in pigs. One such example reported in literature recently where the mixture of boar-unique salivary molecules (Boar Better: androstenone, androstenol and quinoline) induced sexual behaviour in sows after weaning (McGlone et al., 2019). Further, the stimulatory sexual behavioural effect of the mixture (containing all three active molecules) was reported more powerful than any single or binary boar-unique molecule(s). The most biologically-relevant boar pheromone seems to be a mixture of male-unique salivary molecules. Hence, newer identified compounds in SG and saliva can also be used in similar lines. In conclusion, it is shown that seminal gel and saliva can be used effectively to induce estrus in females which can be intensified further in combination with boar saliva.

Identified compounds in chemical forms at standardized doses can be used in healthy pubescent, young or adult boars, experiencing sexual behavioural problems such as low libido as a result of overuse or feeble interest. The identified compounds when applied should be used in conjunction with other alternative collection techniques to improve interest, including - new teaser introduction, creating jealousy, change of environment or the animal handler/collector. It should be kept in mind not to be used on boars, gilts and sows having physical problems incompatible with good utilization of their reproductive potential e.g. overage, acute or chronic back problems, locomotive disorders etc.

## CONCLUSIONS

Mass spectrometry analysis of boar seminal gel, saliva and semen was done to identify the novel compounds having pheromonal properties in pigs. It is found that 9(E)-Octadecenoic acid (Elaidic acid) is the novel pheromonal compound identified in pigs that can be used as biostimulatory enhance reproductive efficiency as well as may be explored further to validate its commercial application in pigs. It can be used an exploit as a nasal spray, ointment or gel for biostimulation purposes. Further, other identified compounds such as tridecenol, undecane, 4-heptane-2-methyl, 2-isopropyl 5-methyl-1-heptanol, hexadecone, 3-ethyl-3 methyl heptone, eicosane, tetracosane etc. may be used in combination as pheromones in pigs. Seminal gel, saliva and semen and their combination can be used to induce oestrus and synchronization in females and training of males for artificial insemination purposes in swine. The combination of seminal gel and saliva has a more intense biostimulation effect than gel alone for the training of males as well as induction of estrus in gilts and sows. Newer commercial preparation with identified compounds may provide a breakthrough for oestrous induction and synchronization in swine reproduction.

## ETHICS STATEMENT

The study was conducted at the Indian Council of Agricultural Research- National Research Centre on Pig, Rani, Guwahati, 781131, India. The experimentation was carried out with prior approval from the Institute Animal Ethics Committee. The approved animal use protocol number was NRCP/CPCSEA/1658|IAEC-54 dated 03^rd^ December, 2019.

## DATA AVAILABILITY STATEMENT

The data in the manuscript is original and included in the manuscript.

## AUTHOR CONTRIBUTIONS

SK, SR, VKG conceived of and designed the experiments, and revised the manuscript. SK,PJD, SB, RI, KB analyzed the data and wrote the manuscript. SK and AC performed the experiments and collected the samples. All authors reviewed and approved the manuscript.

## ACKNOWLEDGMENTS

The authors are grateful to Director, ICAR- National Research Centre on Pig, Rani, Guwahati, 781 131, India for providing the facilities for conduction of the experiment. SK is thankful to Mr. N. Saharia and AR Laboratory staff for helping in the present study.

## SUPPLEMENTAL MATERIAL

Video 1. Representative demonstration for training of male using boar seminal gel rubbed on dummy. Male showing interest in gel and mounts on dummy on exposure to boar seminal gel.

Video 2. Representative demonstration for estrus induction in sow using boar seminal gel kept near to snout of female. Female showing interest in gel for smell, started salivation and champing.

## REFERENCES

Achiraman, S., Archunan, G., Ponmanickam, P., Rameshkumar, K., Kannan, S. and John, G. (2010). 1-Iodo-2 methylundecane [1I2MU]: an estrogen-dependent urinary sex pheromone of female mice. Theriogenolog 74, 345–53. doi: 10.1016/j.theriogenology.2010.01.027. PMID: 20570325

Archunan, G. and Rajanarayanan, S. (2010). Composition for enhancing bull sex libido. Indian Patent No. 244991, Indian Patents Office, India.

Bergman, D. K., Dillwith, J.W., Campbell, R. K. and Eikenbary, R.D. (1990). Cuticular hydrocarbons of the Russian wheat aphid. Southwest. Entomol.15, 91–100. ID=US9044130

Boursnell, J. C, Hartree, E. F. and Briggs, P. A. (1970). Studies on the bulbo-urethral (Cowper’s) gland mucin and seminal gel of the boar. Biochem. J. 117, 981–8. doi: 10.1042/bj1170981

Boursnell, J. C. and Butler, E. J. (1973). Studies on properties of the seminal gel of the boar using natural gel and Gel formed in vitro. J. Reprod. Fertil. 34, 457–63. doi: 10.1530/jrf.0.0340457

Brooks, P. H., and D. J. A. Cole. (1970). The effect of the presence of a boar on the attainment of puberty in gilts. J. Reprod. Fertil. 23, 435–40. doi: 10.1530/jrf.0.0230435

Byers, J. A. (2006). Pheromone component patterns of moth evolution revealed by computer analysis of the Pherolist. J. Anim. Ecol.75, 399–407. doi: 10.1111/j.1365-2656.2006.01060.x

Carlson, D. A., Mayer, M. S., Silhacek, D.L., James, J. D., Beroza, M. and Bierl, B. A. (1971). Sex Attractant Pheromone of the House Fly: Isolation, Identification and Synthesis. Entomology Papers from Other Sources. 11. Science, New Series, Vol. 174, No. 4004 (Oct. 1, 1971), pp. 76–78

Chenoweth, P. J. (1983). Reproductive management procedures in control of breeding. Austral. J. Anim. Prod. 15, 28–33

Choudhary, S. and Kamboj, M.L. (2019).Effect of bull biostimulation on the oestrous behaviour of pubertal Sahiwal (*Bos indicus*) heifers. Anim. Reprod. Sci. 209, 106149. doi: 10.1016/j.anireprosci.2019.106149. PMID: 31514934

Dyck, G.W. (1988). Factors influencing sexual maturation, puberty and reproductive efficiency in the gilt. Canad. J. Anim. Sci., 68, 1–13. doi.org/10.4141/cjas88-001

Dyrmundsson, O. R., and J. L. Lees. (1972). Effect of rams on the onset of breeding activity in Clun Forest ewe lambs. J. Agri. Sci. 79, 269. doi:10.1017/S002185960003224X1972

Farine, J.P., Ferveur, J.F. and Everaerts, C. (2012). Volatile Drosophila cuticular pheromones are affected by social but not sexual experience. PLoS ONE, 7, e40396. doi:10.1371/journal.pone.0040396

Funaro, F. C., Böröczky, K., Vargo, E.L. and Schal, C. (2018). Identification of a queen and king recognition pheromone in the subterranean termite *Reticulitermes flavipes*. Proceedings of the National Academy of Sciences. 115. 201721419. 10.1073/pnas.1721419115. doi: 10.1073/pnas.1721419115

Fujiwara,T. N., Yamagata, N., Takeda, T., Mizunami, M. and Yamaoka, R. (2006). Behavioral responses to the alarm pheromone of the ant *Camponotus obscuripes* (Hymenoptera: Formicidae). Zoological Sci., 23, 353–358. doi.org/10.2108/zsj.23.353

Grant, G.G., Slessor, K.N., Liu, W.M. and Abou-Zaid, M. (2003). (Z,Z)-6,9-heneicosadien-11-one, labile sex pheromone of the white marked tussock moth, *Orgyia leucostigma*. J. Chem. Ecol. 29, 589–601. doi: 10.1023/a:1022802821338

Karthikeyan, K., Muniasamy, S., Ganesh, D. S., Achiraman, S., Saravanakumar, V. R. and Archunan, G. (2013). Faecal chemical cues in water buffalo that facilitate estrus detection. Anim. Reprod. Sci. 138, 163–67. doi.org/10.1016/j.anireprosci.2013.02.017

Karthikeyan, K. and Archunan. G. (2013). Gas chromatographic mass spectrometric analysis of estrus specific volatile compounds in buffalo vaginal mucus after initial sexual foreplay. J. Buffalo Sci., 1–7. doi.org/10.6000/1927-520x.2013.02.01.1

Kekan, P.M., Ingole, S.D., Sirsat, S.D., Bharucha, S.V., Kharde S.D. and Nagvekar, A.S. (2017). The role of pheromones in farm animals - A review. Agri. Rev. 38, 83–93. doi: 10.18805/ag.v38i02.7939

Kirkwood, R. N., Forbes, J. M. and Hughes, P. E. (1981). Influence of boar contact on attainment of puberty in gilts after removal of the olfactory bulbs. J. Reprod. Ferti. 61,193–6. doi: 10.1530/jrf.0.0610193

Liu, B., Syu, K.J., Zhang, Y.X., Gupta, S., Shen, Y.J., Chien, W.J. et. al. (2020). Practical synthesis and field application of the synthetic sex pheromone of rice stem borer. *Chilo suppressalis* (Lepidoptera: Pyralidae). J. Chemistry. Article ID 5632534, 9 pages. doi.org/10.1155/2020/5632534

McGlone, J., Devaraj, S. and Garcia, A. (2019). A novel boar pheromone mixture induces sow estrus behaviors and reproductive success. Appl. Anim. Behav. Sci. 219. 104832. doi.org/10.1016/j.applanim.2019.104832

Meinwald, J., Boriack, C.J., Schneider, D., Boppre, M., Wood, W.F. and Eisner, T. (1974). Volatile ketones in the hair pencil secretion of danaid butterflies (Amauris and Danaus). Experientia 32,721–722. doi:10.1007/BF01924148

Melrose, D.R., Reed, H.C. and Patterson, R.L., 1971. Androgen steroids associated with boar odour as an aid to the detection of oestrus in pig artificial insemination. British Vet. J. 127, 497–502. doi: 10.1016/s0007-1935(17)37337-2

Mendki, M.J., Khetiswari, G., Prakash, S., Suryanarayana, M. Malhotra, R.C., Rao, K.M. et al. (2000). Heneicosane: An oviposition-attractant pheromone of larval origin in Aedes aegypti mosquito. Curr. Sci. 78, 1295–1296. Corpus ID: 82715500

Minaeimoghadam, M., Askarianzadeh, A., Imani, S., Shojaei, M., Larijani K. and Abbasipour H.(2017): Identification of chemical compounds of the pheromone in different ages of female adults of the clearwing moth, *Paranthrene diaphana* Dalla Torre and Strand, Arch. Phytopathol. Plant Prot. 50, 1019–1033. doi: 10.1080/03235408.2017.1411174

Mitaka, Y., Matsuyama, S., Mizumoto, N., Matsuura K. and Akino T. (2020). Chemical identification of an aggregation pheromone in the termite *Reticulitermes speratus*. Sci. Rep. 10, 7424 doi.org/10.1038/s41598-020-64388-4

Mittersteiner, M., Linshalm, B.L., Vieira, A.P.F., Brondani, P.B., Scharf, D.R. and de Jesus, P.C. (2017). Convenient enzymatic resolution of (R,S)-2-methylbutyric acid catalyzed by immobilized lipases. Chirality 30, 106–111. doi: 10.1002/chir.22779

Mizunami, M., Yamagata N. and Nishino H. (2010). Alarm pheromone processing in the ant brain: an evolutionary perspective. Front. Behav. Neurosci. 8, 4:28. doi: 10.3389/fnbeh.2010.00028. PMID: 20676235; PMCID: PMC2912167

Molander, M., Jimmy H., Inis W., Jocelyn M., and Mattias, L. (2019). The male-produced aggregation-sex pheromone of the cerambycid beetle *Plagionotus detritus* ssp. detritus. J. Chem. Ecol. 45. doi: 10.1007/s10886-018-1031-4

Oldham, C. M., G. B. Martin, and T. W. Knight. (1978). Stimulation of seasonally anovular ewes by rams. Time from introduction of rams to the preovulatory LH surge and ovulation. Anim. Reprod.Sci. 1, 283–290., doi:283-290. 10.1016/0378-4320(79)90013-7

Pearce, G. P. and Oldham, C. M. (1988). Importance of non-olfactory ram stimuli in mediating ram-induced ovulation in the ewe. J. Reprod. Fert. 84:333. doi.org/10.1530/jrf.0.0840333

Pickett, J. A., Williams, I. H. and Martin, A. P. (1982). (Z)-11-eicosen-1-ol, an important new pheromonal component from the sting of the honey bee, *Apis mellifera*, L. J. Chem. Ecol. 8, 163–175. doi: 10.1007/BF00984013

Rajanarayanan, S. and Archunan, G. (2011). Identification of urinary sex pheromones in female buffaloes and their influence on bull reproductive behavior. Res. Vet. Sci. 91, 301–305. doi: 10.1016/j.rvsc.2010.12.005

Ray, A.M., Swift, I.P., Moreira, J.A., Millar, J.G. and Hanks, L.M. (2009). (R)-3-hydroxyhexan-2-one is a major pheromone component of *Anelaphus inflaticollis* (Coleoptera: Cerambycidae. Environ. Entomol. 38,1462–1466. doi: 10.1603/022.038.0514.

Ray, A.M., Barbour, J.D., McElfresh, J.S., Moreira, J. A., Swift, I., Wright, I. M. et al. (2012). 2,3-Hexanediols as sex attractants and a female-produced sex pheromone for cerambycid beetles in the prionine genus tragosoma. J. Chem. Ecol. 38, 1151–1158 doi /10.1007/s10886-012-0181-z

Rekwot, P.I.Ogwu, D. and Oyedipe, E.O. (2000). Influence of bull biostimulation, season and parity on resumption of ovarian activity of zebu (*Bos indicus*) cattle following parturition. Anim. Reprod. Sci., 63(1-2),1–11. doi: 10.1016/s0378-4320(00)00163-9. PMID: 10967236

Rekwot, P.I., Ogwub, D., Oyedipe E.O. and Sekonia, V.O. (2001). The role of pheromones and biostimulation in animal reproduction. Anim. Reprod. Sci., 65, 157–170. doi.org/10.1016/S0378-4320(00)00223-2

Renou, M., Descoins, C., Priesner, E.Gallois, M. and Lettere, M. (1981). A study of the sex pheromone of the leek moth, *Acrolepiopsis assectella* (Lepidoptera: Acrolepiidae). Entomol. Exp. Appl. 29,198–208

Rowlinson, P., and Bryant. M. J. (1974). Rebreeding sows during lactation - A system for overcoming lactational anestrus with reference to the effect of the male. III^rd^ International Pig Veterinary Congress. 5:1

Shelton, M. (1960). Influence of the presence of a male goat on the initiation of oestrous cycling and ovulation of Angora does. J. Anim. Sci.19,368–375

Takahashi, G., Maeda, M., Kimura. Y. and Funahashi. H. (2014).Biochemical analysis of component in seminal gel secreted with boar semen. Reprod. Fertil. Dev. 27, 100–101. doi.org/10.1071/RDv27n1Ab16

Thibout, E., Ferary, S. and Auger, J. (1994). Nature and role of sexual pheromones emitted by males of *Acrolepiopsis assectella* (LEP.). J. Chem. Ecol. 20, 1571–81.doi: 10.1007/BF02059881

Thomson, L.H. and Savege, J.S. (1978). Age at puberty and ovulation rate in gilts in confinement as influenced by exposure to a boar. J. Anim. Sci.,47, 1141–1146. doi: 10.2527/jas1978.4751141x

Varner, E., Gries, R., Takács, S., Fan, S. and Gries, G. (2019). Identification and field testing of volatile components in the sex attractant pheromone blend of female house mice. J. Chem. Ecol. 45, 18–27. doi: 10.1007/s10886-018-1032-3

Wattanachaiyingcharoen, W., Phanmuangma, W., Boonphong, S., Suphrom, N., and Prasanpan, S. (2020). Sex pheromone and pattern of mating communication of fireflies in subfamily lampyrinae (coleoptera: lampyridae). PSRU J. Sci. Tech.5, 35–46

Wikimedia. Heneicosane. (2014). https://commons.wikimedia.org/wiki/File:Heneicosane_2.JPG [Accessed April 08, 2022]

Wittko, F. (1999). In Comprehensive Natural Products Chemistry. Miscellaneous Natural Products Including Marine Natural Products, Pheromones, Plant Hormones, and Aspects of Ecology.

Würf, J., Pokorny, T., Wittbrodt, J., Millar, J.G. and Ruther, J. (2020). Cuticular hydrocarbons as contact sex pheromone in the parasitoid wasp urolepis rufipes. Front. Ecol. Evol. 8, 180.doi: 10.3389/fevo.2020.00180

Yadav, A.K. and Tissot, P. (1984). Electrochemical synthesis of the sex attractant pheromone of the housefly *Musca domestica* (z)-9-tricosene. Helv. Chim. Acta. 67, 1698–1701. doi: 10.1002/hlca.19840670705

